# Osthole Suppresses Prostate Cancer Progression by Modulating PRLR and the JAK2/STAT3 Signaling Axis

**DOI:** 10.1101/2024.12.30.630802

**Authors:** Linjun Yan, Yuanqiao He, Qi Cui, Xiaohong Wang, Feng Lv, Keyue Cao, Yuanjian Shao

**Author notes:** Correspondence: Yuanjian Shao. First author: Linjun Yan.

## Abstract

Prostate cancer is a common malignancy in men with limited effective treatment options, highlighting the urgent need for novel therapeutic approaches. Osthole, a natural coumarin compound with antitumor properties, has demonstrated potential in targeting various cancers. This study investigates the therapeutic effects of Osthole in prostate cancer by focusing on its ability to target the Prolactin Receptor (PRLR) and modulate the JAK2/STAT3 signaling pathway. Using a combination of network pharmacology, in vitro assays, and in vivo experiments, we first employed network pharmacology to predict Osthole’s potential targets, identifying 68 targets shared with prostate cancer, including AKT1, TNF, IL6, STAT3, and CTNNB1. We then confirmed these targets and assessed the effects of Osthole on cell proliferation, migration, and apoptosis using the Cell Counting Kit-8 (CCK-8) and transwell invasion assays. To further elucidate the molecular interactions and protein expression levels, we employed molecular docking and western blot analysis. Our findings revealed that Osthole significantly inhibited prostate cancer cell proliferation and migration in a dose-dependent manner and reduced tumor volume in in vivo assays. Western blot analysis also indicated that Osthole downregulated PRLR expression and decreased the phosphorylation of JAK2 and STAT3, suggesting the inhibition of the JAK2/STAT3 pathway. These results collectively highlight the therapeutic potential of Osthole in targeting prostate cancer cells through PRLR and modulating the JAK2/STAT3 pathway, warranting further clinical exploration.

## 1. Introduction

Prostate cancer, which originates in the prostate gland in males, is recognized as one of the most prevalent malignancies within the male urogenital system[1]. Globally, prostate cancer ranks third in occurrence and sixth in mortality among men[2]. Prostate cancer is predominantly treated with surgical resection, castration, and radiochemotherapy[3]. However, owing to the limited effectiveness of conventional therapeutic approaches in improving the survival of patients with poor prognoses, there is an urgent need to develop novel treatment modalities. Cnidium monnieri extract, also known as Osthole, is a naturally occurring compound found in various medicinal plants, particularly in species such as Cnidium monnieri and Angelica sinensis. This compound, classified within the coumarin class of chemical compounds, exhibits a diverse array of biological activities, including antitumor, anti-inflammatory, neuroprotective, osteogenic, cardiovascular protective, antimicrobial, and antiparasitic effects[4]. There is documented evidence in the literature that Osthole has been utilized for the treatment of various cancers, including gastric, bladder, and breast cancers[5–7]. Additionally, only a limited number of studies have documented the inhibitory effects of Osthole on prostate cancer in vitro[8]. However, research on its in vivo effects remains limited.

The prolactin receptor (PRLR), a key member of the growth hormone receptor family, has been closely associated with hormone-dependent cancers such as breast and ovarian cancers [9]. However, its specific role and underlying mechanisms in prostate cancer remain poorly understood. PRLR is known to activate multiple downstream signaling pathways, among which the JAK2/STAT3 pathway plays a crucial role in tumor cell proliferation, immune evasion, and therapeutic resistance [10]. Investigating the regulatory mechanisms of PRLR and its downstream JAK2/STAT3 signaling axis in prostate cancer may provide a scientific foundation for the development of novel targeted therapies. Previous studies have demonstrated that Osthole can regulate pathways such as PI3K/Akt and MAPK, thereby inhibiting tumor growth in gastric and bladder cancers [11]. However, whether Osthole exerts its anticancer effects in prostate cancer by modulating PRLR and the JAK2/STAT3 signaling axis has not been thoroughly investigated.

Therefore, in this study, we utilized network pharmacology to predict the common targets of Osthole and prostate cancer, which were subsequently validated through a series of in vivo and in vitro experiments. This study could potentially contribute to the development of novel therapeutic agents for the treatment of prostate cancer.

## 2. Materials and Methods

### 2.1 Experimental Materials

Prostate cancer cell lines RM1, 22RV1, PC-3, and DU145, along with RPMI 1640 culture medium and other reagents, were procured from Fuheng Biological Technology Co., Ltd. Osthole was obtained from Luoen Biological Technology Co., Ltd., and the RPMI 1640 culture medium was acquired from Tianjin Haoyang Biological Products Technology Co., Ltd.

### 2.2 Network pharmacological analysis

#### 2.2.1 Selection of Osthole Targets

Three-dimensional structural information of the compound was retrieved from the PubChem database (https://pubchem.ncbi.nlm.nih.gov/). Target proteins of the compound were identified using the Swiss Target Prediction (STP) database (http://www.swisstargetprediction.ch/) with a probability threshold greater than 0. Additionally, the BATMAN-TCM2.0 database (http://bionet.ncpsb.org/batman-tcm/) was used to filter target proteins with a cutoff value exceeding 0.85. The resulting target proteins were consolidated by eliminating duplicates to yield a list of potential target proteins.

#### 2.2.2 Collection of Prostate Cancer Targets

Genes linked to prostatic cancer were identified by searching the term ‘prostatic cancer’ in DisGeNET(https://www.disgenet.org/), Genecards(https://www.genecards.org/), and OMIM databases(https://omim.org/) with specific filters: a GDA score >0.1 DisGeNET, a relevance score >10 in Genecards, and relevant gene entries in OMIM. After removing duplicates, the compiled gene list was intersected with the target proteins of the compound to identify the shared targets.

#### 2.2.3 Construction of Protein-protein interaction (PPI)

The intersecting target dataset was imported into the STRING database (https://string-db.org/) with Homo sapiens selected as the species, and the minimum required interaction score was set to 0.4. The ‘string_interactions_short.tsv’ file was then downloaded and imported into Cytoscape 3.7.2 for network visualization. Network topology analysis was performed using the CytoNCA plugin.

#### 2.2.4 Construction of Drug-Compound-Target Network Diagram

The intersection of disease and compound targets identified a set of genes potentially responsible for therapeutic effects. Data corresponding to these genes were extracted from the drug database and compiled into a network, which was then saved as ‘network.xlsx’. Subsequently, this network data file was imported into the Cytoscape 3.7.2 to visualize the network.

#### 2.2.5 Pathway Analysis of Targets

Enrichment analysis for the intersecting targets was performed using the R packages ‘ClusterProfiler’, ‘org.Hs.eg.Db’, and ‘ggplot2’. Gene annotation was sourced from the ‘package. The analysis utilized an adjusted p-value threshold of 0.05 and a q-value threshold of 0.05 to determine statistical significance.

#### 2.2.6 Molecular Docking

Top-ranked macromolecules were docked with small molecules from the PubChem database using protein structures from the AlphaFold Database. Protein structures were prepared using AutoDockTools 5.6, and small molecules were processed using Open Babel and AutoDock. Docking was performed using AutoDock, and the results were visualized using PyMOL.

### 2.3 Cell Functional Assays

#### 2.3.1 Cell Proliferation Assay

22RV1, PC-3, DU145, and RM1 cell lines were cultured in RPMI-1640 medium supplemented with 10% FBS. The cells were then incubated in a humidified incubator at 37°C with 5% CO2. Cells in the logarithmic growth phase at approximately 90% confluence were treated with trypsin, seeded at 10,000 cells per well in a 96-well plate, and cultured for 24 hours. Subsequently, the cells were treated with Osthole or Saline and incubated for 48 hours. After treatment, CCK-8 was added, and the cells were incubated for 1-4 hours. Absorbance was measured at 450 nm using an enzyme-linked microplate reader. Each experiment was performed in triplicate, and the experiment was repeated at least twice. Data are presented as mean ± standard deviation.

#### 2.3.2 Transwell Invasion Assay

Matrigel (BD Biosciences, San Jose, CA, USA) was diluted 1:3 in serum-free RPMI-1640, and 50 μL was used to pre-coat the Transwell inserts. After 48 hours of Osthole treatment, RM1 cells were collected and suspended in serum-free DMEM at a density of 1×10^5 cells/mL, and 500 μL of this cell suspension was added to the upper chamber. Next, 600 μL of RPMI-1640 containing 10% FBS was added to the lower chamber. RM1 cells were incubated for 24 hours at 37°C with 5% CO2 to allow for invasion. Invasive cells were fixed with 4% paraformaldehyde for 15 minutes at room temperature and then stained with 0.1% crystal violet for 10 minutes at room temperature. Finally, the invading cells were photographed and counted under an inverted microscope, with five random non-overlapping fields selected for cell counting.

### 2.4 Tumor Inoculation and Monitoring

RM1 and 22RV1 cell suspensions, each containing 1×10^7^ cells in 0.1 mL PBS, were injected subcutaneously into the right hind limb of nude mice to induce tumor growth. Tumor volume was monitored and measured biweekly with calipers, allowing the tumors to reach an approximate size of 100 mm^3^ before proceeding with experimental manipulation. Once the tumors reached 100 mm^3^, the mice were randomly assigned to two groups, each consisting of five animals: a treatment group and a control group. The treatment group received injections of Osthole at a dosage of 200 mg/kg via the intraperitoneal route, administered daily for three weeks (200 mg/kg, IP, once a week for three weeks). In contrast, the control group was injected with an equal volume of saline.

Tumor volume and weight were measured twice weekly, with observations of tumor morphology and ulceration status before and after each treatment. Tumor volume was expressed in cubic millimeters (mm^3^), calculated using the formula: (V = 0.5×a×b^2^), where a and b represent the longest and shortest diameters of the tumor, respectively. The experimental endpoint was determined based on four criteria: 1) termination of the experiment after 21 days of drug intervention if the average volume of the model control group exceeded 500 mm^3^; otherwise, the intervention period was extended; 2) a tumor volume exceeding 2000 mm^3^ in the mice; 3) ulceration and necrosis of the mouse tumor; and 4) weight loss exceeding 20% of the normal animal weight. At the end of the experiment, the tumors from the mice were excised and weighed to obtain the tumor mass.

### 2.4 Bioinformatics Analysis of PRLR, JAK2, and STAT3 in Prostate Cancer Using TCGA Data

RNA-seq data and clinical information for prostate cancer patients were obtained from the TCGA-PRAD dataset, including 496 tumor samples and 52 normal samples. Differential gene expression analysis was performed using the ‘limm’ R package, with |log2FC| ≥ 1 and adjusted p-value < 0.01 as significance thresholds. Kaplan-Meier survival analysis was conducted for PRLR, JAK2, and STAT3 using the ‘survival’ and ‘survminer’ R packages, categorizing patients into high– and low-expression groups based on the median expression value, and survival differences were assessed via log-rank tests. Gene Ontology (GO) and KEGG pathway enrichment analyses of differentially expressed genes were performed using ‘clusterProfiler‘, with adjusted p-value < 0.05 as the cutoff. Gene Set Enrichment Analysis (GSEA) for PRLR, JAK2, and STAT3 was conducted with the ‘fgse’ and ‘enrichplot’ R packages to explore associated pathways. All analyses and visualizations were performed in R (version 4.2.0) with additional packages including ‘ggplot2‘, ‘pheatmap‘, and ‘AnnotationDbì.

### 2.5 Western Blot Analysis of RM1 Cells

RM1 cells were lysed for protein extraction and equal amounts of protein were separated by SDS-PAGE and transferred to PVDF membranes. After blocking, the membranes were incubated with primary antibodies against JAK2, p-JAK2, STAT3, p-STAT3, and β-actin overnight at 4°C, followed by incubation with HRP-conjugated secondary antibodies. Protein bands were visualized using Enhanced Chemiluminescence, quantified by densitometry with ImageJ, and normalized to β-actin.

### 2.6 Statistical Analysis

All calculations were performed using Prism 6, and the data are presented as mean ± standard deviation (SD). Multiple group comparisons were made using two-way ANOVA, with a p-value of less than 0.05 considered statistically significant.

## 3. Results

### 3.1 Network Pharmacology Analysis of Osthole Targets and Pathways in Prostate Cancer

Potential targets of Osthole were predicted using the Swiss Target Prediction and BATMAN-TCM2.0 databases, and 68 valid targets were identified after excluding the ineffective components. Concurrently, 2507 prostate cancer-related targets were filtered from the DisGeNET, Genecards, and OMIM databases. Venn diagram analysis revealed 68 common targets between Osthole and prostate cancer (Figure 1A). Subsequently, PPI network analysis was conducted on these targets using the STRING database, which suggested that genes such as AKT1, TNF, IL6, STAT3, and CTNNB1 might be key targets for Osthole in the treatment of prostate cancer (Figure 1B-C).

**Fig 1.**
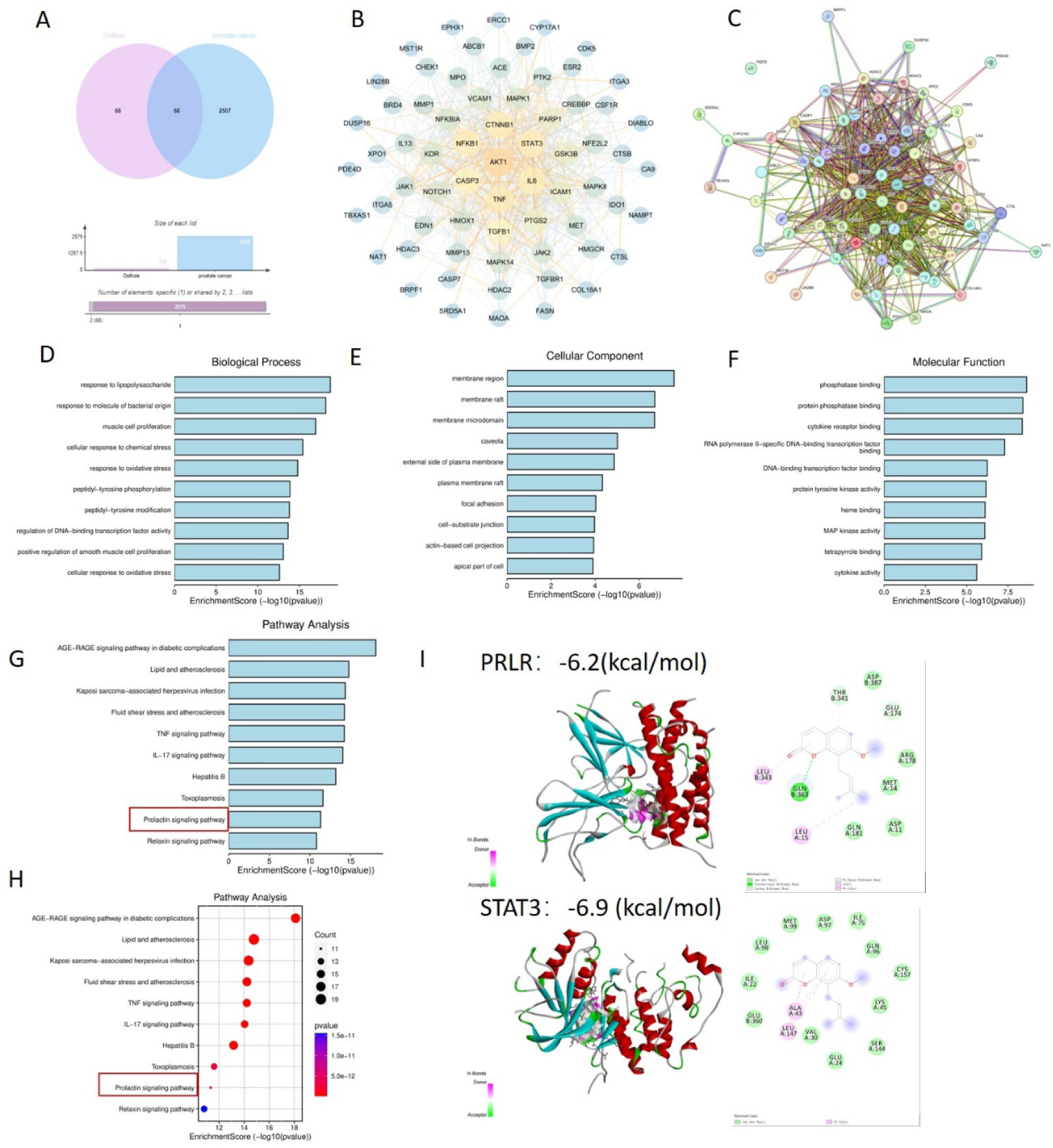
Network Pharmacology Analysis. A and B: Analysis of databases revealed 68 shared targets for Osthole in prostate cancer. C: PPI network analysis using STRING database identified AKT1, TNF, IL6, STAT3, and CTNNB1 as possible therapeutic targets of Osthole in prostate cancer. D to F: Intersection and GO analysis revealed Osthole’s association with critical prostate cancer-related pathways and functions. G and H: Kyoto Encyclopedia of Genes and Genomes (KEGG) analysis revealed the involvement of Osthole in the PRL pathway and other significant cancer-related pathways. I: STAT3 and PRLR displayed strong binding to Osthole, indicating likely therapeutic targets.

Furthermore, an intersection approach was employed to identify disease-related targets from the drug data, which were then analyzed using Cytoscape 3.7.2. Gene Ontology (GO) enrichment analysis yielded a total of 3993 results, including 3469 biological process (BP) terms primarily associated with lipopolysaccharide, response to molecules of bacterial origin, and muscle cell proliferation; 191 cellular component (CC) terms related to membrane regions, membrane rafts, and membrane microdomains; and 371 molecular function (MF) terms involving phosphatase binding, protein phosphatase binding, and cytokine receptor binding (Figure 1D-F). The Kyoto Encyclopedia of Genes and Genomes (KEGG) enrichment analysis identified 226 pathways, predominantly involving the advanced glycation end-product receptor for advanced glycation end-product signaling pathway in diabetic complications and lipid and atherosclerosis, suggesting that Osthole may intervene in prostate cancer through these pathways (P < 0.05)(Figure 1G-H).

Additionally, molecular docking with a binding energy of less than –1 kcal/mol indicates active binding, while a value less than –5 kcal/mol suggests a stronger binding affinity. The molecular docking results demonstrated that STAT3 and PRLR exhibit favorable binding activities with cisplatin, with binding energies of –6.9 and –6.2 kcal/mol, respectively, indicating a strong affinity of these proteins for Osthole (Figure 1I).

### 3.2 Effects of Osthole on Prostate Cancer Cell Proliferation, Migration, and Apoptosis

We evaluated the effect of Osthole on the proliferation of prostate cancer cell lines using the CCK-8 cell proliferation assay (Figure 2A). After treating the cells with 0, 50, 100, 150, 200, 300, 400 and 500 μM of Osthole for 48 hours, we found that it exerted significant cytotoxicity on the 22RV1 and PC-3 cell lines, the half-maximal inhibitory concentration (IC50) values for the prostate cancer cell lines DU145, PC-3, RM1, and 22RV1 were determined to be 66.44 μM, 238 μM, 102 μM, and 319.5 μM, respectively. These values indicate that the proliferation of the respective cell lines was inhibited by 50%. Compared with the control group, cisplatin-treated cells showed significant suppression of proliferation at all concentrations tested (P<0.05), with cell viability decreasing as Osthole concentration increased, indicating a dose-dependent inhibitory effect on prostate cancer cell proliferation.

**Fig 2.**
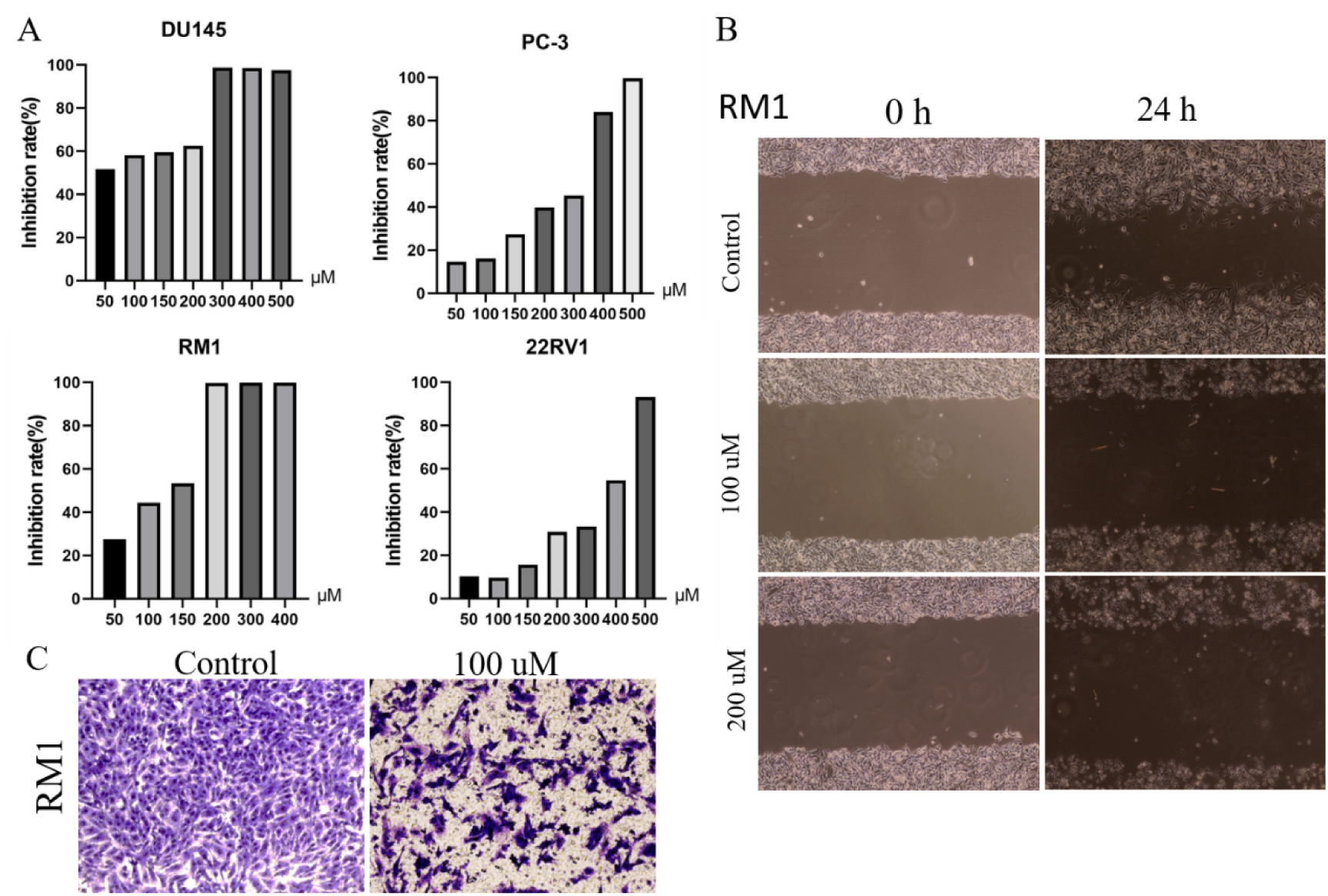
Cell Function Analysis. A)Osthole exhibited cytotoxic effects on the prostate cancer cell lines 22RV1, PC-3, DU145, and RM1 in a concentration-dependent manner, with IC50 values of 66.44, 238, 102, and 319.5 µM, respectively. B) The effect of Osthole on the migration of RM1 prostate cancer cells was assessed using a scratch assay. RM1 cells were cultured to confluence, and a uniform scratch was created across the cell monolayer. After treating the cells with various concentrations of Osthole for 24 hours, the migration of cells into the scratch area was observed. C) The invasiveness of RM1 prostate cancer cells was evaluated using a Transwell invasion assay. RM1 cells were seeded in the upper chamber of a Transwell insert coated with Matrigel and treated with Osthole at different concentrations for 48 hours. The number of cells that migrated through the Matrigel membrane to the lower chamber was quantified.

Additionally, the effect of Osthole on the migratory ability of prostate cancer cells was assessed using the scratch assay (Figure 2B). The results indicated that Osthole-treated groups had significantly reduced cell migration distance 24 h post-scratch (P<0.05), suggesting that Osthole effectively inhibits cell migration.

The Control group showed a high number of cells that invaded the Matrigel-coated membrane, indicating a strong invasive capability. The cells were densely packed and uniformly distributed, suggesting their active migration and invasion. The Osthole group showed a marked decrease in the number of invading cells. The cells appeared less dense and more scattered(Figure 2C), indicating that Osthole significantly impaired their invasive ability. This reduction in cell invasion suggests that Osthole effectively inhibits the cellular mechanisms involved in invasion and metastasis.

### 3.3 In Vivo Tumor Growth Inhibition by Osthole in Prostate Cancer

For this study, we selected the human prostate cancer cell line 22RV1 and murine prostate cancer cell line RM1 to establish a CDX model for investigating prostate cancer progression. Given the most pronounced sensitivity of the human-derived 22RV1 cell line to Osthole, this cell line was selected for subsequent experiments. Additionally, the murine RM1 cell line was used to eliminate the potential confounding effects of spontaneous tumorigenesis in mice on the experimental outcomes. Osthole treatment significantly suppressed tumor growth in mice. Compared with the untreated control group, mice administered Osthole showed a markedly reduced rate of tumor volume expansion during the treatment period (P < 0.05)(Figure 3A,6A). Upon completion of the treatment regimen, the average tumor size in RM1 tumor-bearing mice treated with Osthole was significantly reduced compared to that in the control group (P < 0.01) (Figures 3B-C). Although a decreasing trend in tumor volume was observed in 22RV1 tumor-bearing mice subjected to Osthole treatment, the reduction in tumor weight was not statistically significant (Figures 4B-C). This discrepancy may be attributed to the limited number of mice in the study, which could have resulted in insufficient power to detect significant differences.

**Fig 3.**
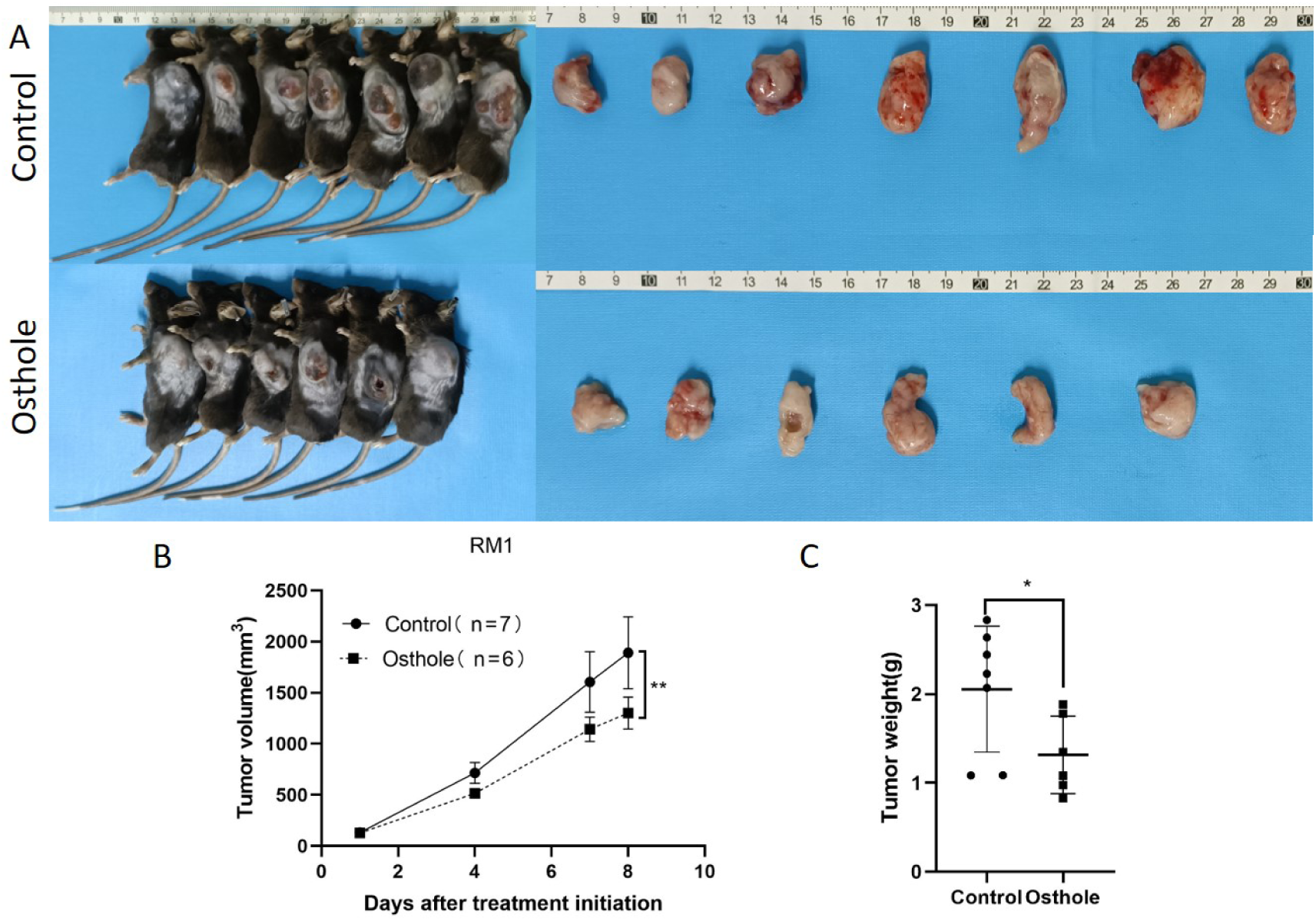
Tumor Growth and Weight Analysis in RM1-Bearing Mice. A) Tumor entities from RM1-bearing mice at day 8 are shown for both control and Osthole-treated groups. B)The subsequent growth curve documented the progressive increase in tumor volume over time (P < 0.01). C)A bar graph quantified the tumor weights, yielding a comparative assessment of the neoplastic burden in RM1 tumor-bearing mice (P < 0.05).

**Fig 4.**
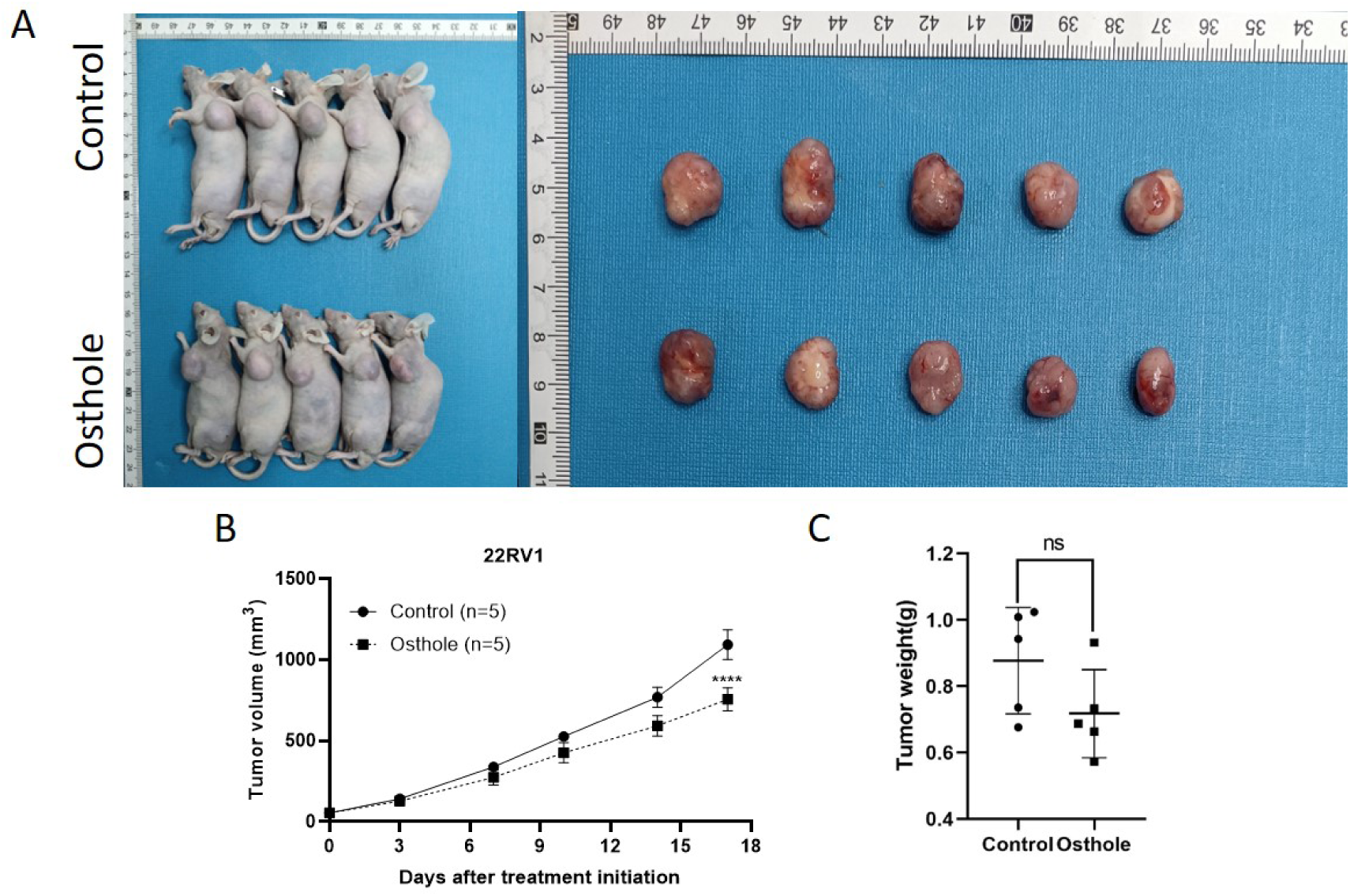
Tumor Growth and Weight Analysis in 22RV1-Bearing Mice. A)Tumor entities from 22RV1-bearing mice at day 17 are presented for both the control and Osthole-treated groups. B)The growth curve on day 17 demonstrated a statistically significant increase in tumor volume over time in the control group (P < 0.0001). C)A bar graph quantified the tumor weights, providing a comparative assessment of the neoplastic burden in 22RV1 tumor-bearing mice.

**Fig 5.**
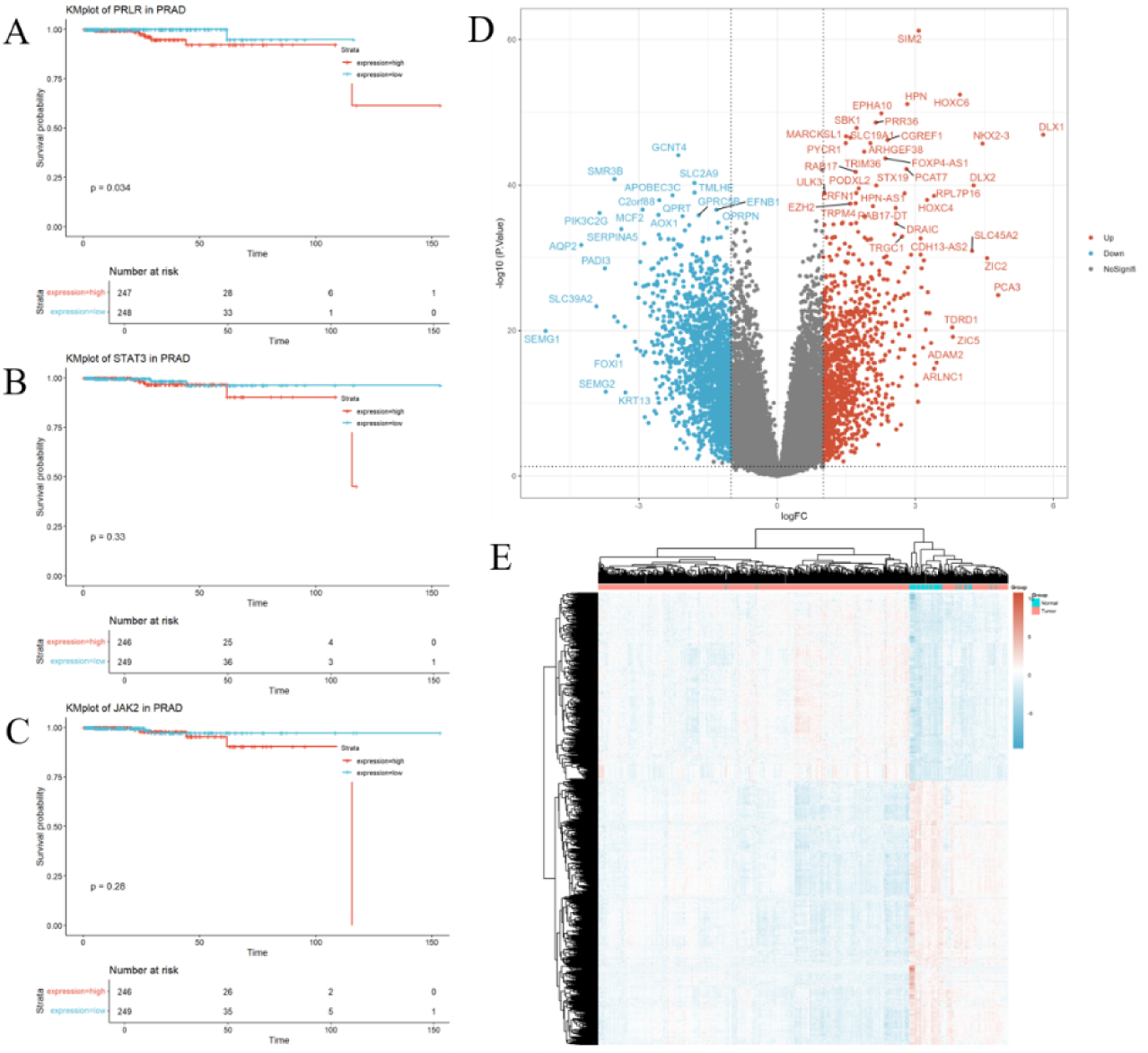
Correlation of PRLR, JAK2, and STAT3 genes with prognosis. (A–C) Kaplan-Meier survival curves for PRLR, JAK2, and STAT3 genes in prostate cancer patients. Survival curves for the high-expression group (red) and the low-expression group (blue) show distinct trends. High expression of the PRLR gene is significantly associated with worse survival outcomes (p = 0.034), whereas the expression levels of JAK2 and STAT3 genes are not significantly correlated with survival (p = 0.28 and p = 0.33, respectively). (D) Volcano plot illustrating the expression of differentially expressed genes in prostate cancer. The x-axis (logFC) denotes the log-fold change in gene expression, and the y-axis (-log10(P.Value)) denotes the negative log-transformed P-value. Red dots indicate the 1,087 significantly upregulated genes, blue dots indicate the 1,829 significantly downregulated genes, and gray dots represent the remaining genes that do not show significant changes. Out of a total of 32,151 genes analyzed, 30,235 genes are not significantly differentially expressed. (E) Heatmap illustrating the expression patterns of differentially expressed genes between disease and normal samples. Rows represent genes, columns represent samples, and the color gradient from blue to red corresponds to gene expression levels ranging from low to high.

**Fig 6.**
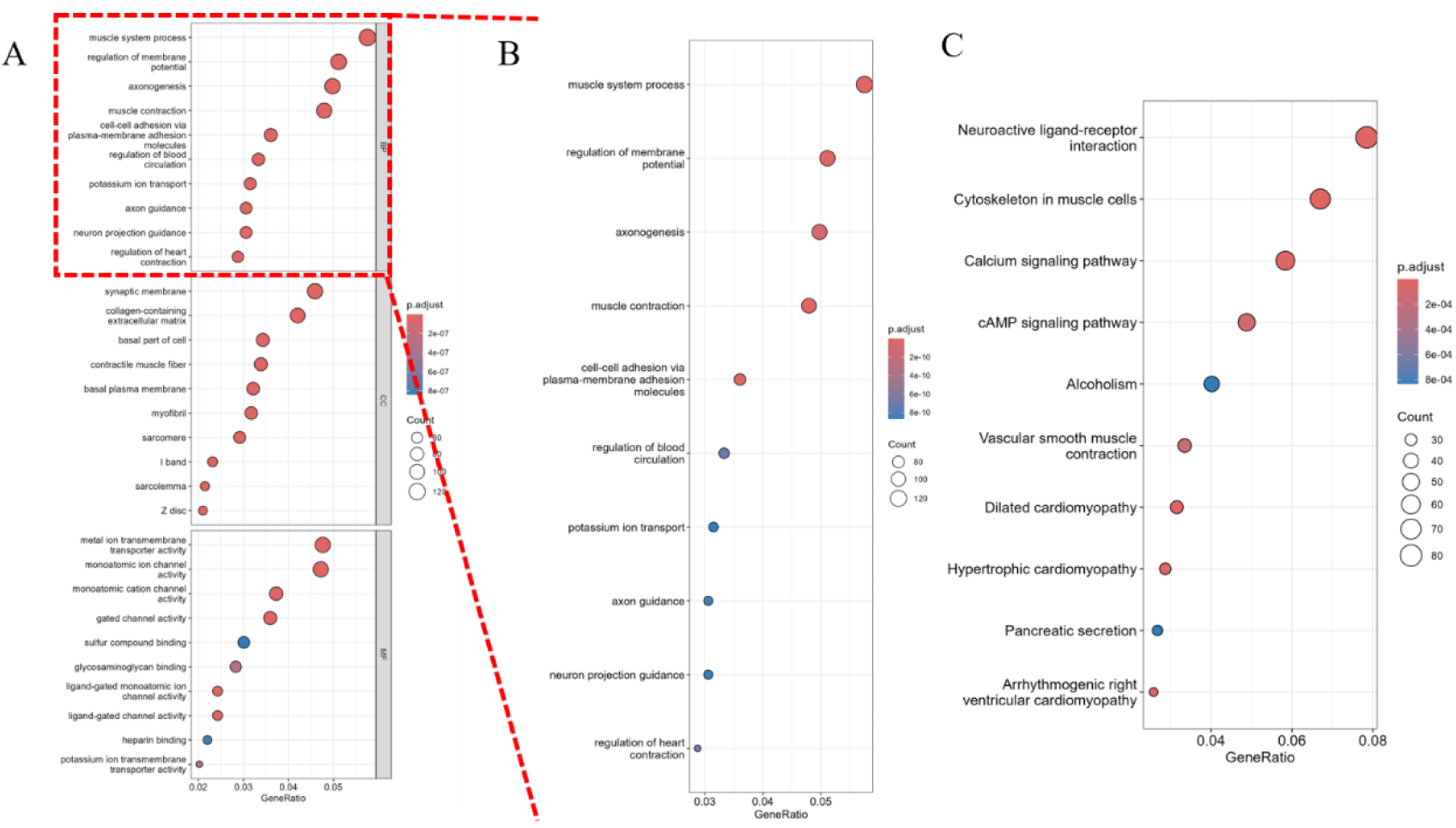
Pathway analysis of differentially expressed genes. (A) Gene Ontology (GO) analysis, including enrichment analysis of biological processes (BP), molecular functions (MF), and cellular components (CC). The x-axis represents the gene ratio (GeneRatio), and the y-axis lists the significantly enriched pathways. The size of the dots represents the gene count (Count), and the color intensity indicates the significance of the adjusted p-value (p.adjust), with darker colors representing smaller p.adjust values and higher significance. (B) KEGG pathway analysis. The x-axis represents the gene ratio (GeneRatio), and the y-axis lists the significantly enriched pathways. The size of the dots represents the gene count (Count), and the color intensity indicates the significance of the adjusted p-value (p.adjust).

### 3.4 Correlation Between PRLR and Prognosis in Prostate Cancer

To investigate the correlation between PRLR and prostate cancer, we identified differentially expressed genes between prostate cancer patients and normal individuals from public databases. Bioinformatics analysis was then performed to assess the relationship between the expression levels of PRLR, JAK2, and STAT3 genes and the survival of prostate cancer patients. The results showed that high expression of the PRLR gene was significantly associated with poor survival outcomes, while JAK2 and STAT3 gene expression levels were not correlated with prognosis (Fig. 5A–C). The volcano plot indicated significant upregulation of genes such as SIM2, HPN, and HOXC6 in prostate adenocarcinoma (PRAD), and significant downregulation of genes such as SLC39A2, SLC39A6, and SLC39A10 (Fig. 5D). A heatmap further demonstrated the expression patterns of differentially expressed genes in prostate cancer (Fig. 5E).

Subsequently, the differentially expressed genes were subjected to GO and KEGG enrichment analyses. Significant biological processes included muscle system processes, regulation of membrane potential, and axon guidance; significant molecular functions involved regulation of cardiac contraction, neuron projection guidance, and regulation of blood circulation; and significant pathways included neuroactive ligand-receptor interaction, muscle cytoskeleton, and calcium signaling pathway (Fig. 6). These findings highlight the potential roles of the gene sets in biological functions and disease processes.

Finally, we conducted enrichment analysis of pathways associated with PRLR, JAK2, and STAT3 genes. The results revealed significant enrichment of gene sets related to muscle system processes, regulation of membrane potential, and axonogenesis under disease conditions (Fig. 7).

**Fig 7.**
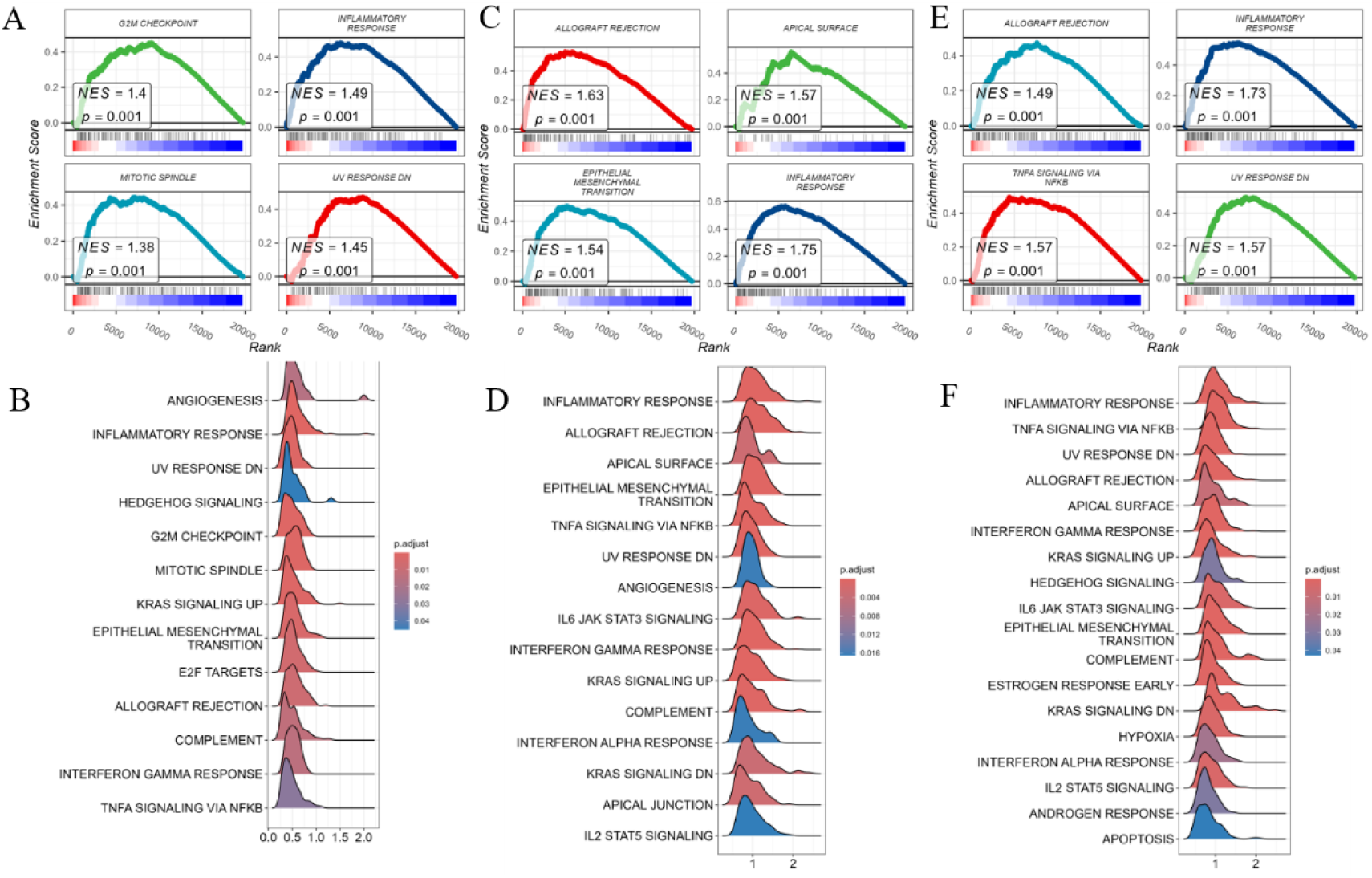
Gene set enrichment analysis (GSEA) of PRLR, STAT3, and JAK2. (A–C) Enrichment analysis of pathways associated with PRLR, JAK2, and STAT3 genes, respectively. Red and blue bar plots represent downregulated (DN) and upregulated (UP) gene sets, respectively. The closer the normalized enrichment score (NES) is to 1, the higher the enrichment level of the gene set in the samples. The enrichment score (GeneRatio) and the adjusted p-value (p.adjust) are indicated by the size and color of the dots, respectively.

### 3.5 JAK2/STAT3 Pathway Regulation by Osthole Through PRLR Interaction

To uncover the molecular underpinnings of the impact of Osthole on prostate cancer, this study utilized a western blot assay to methodically evaluate the expression levels of PRLR, as well as the expression and phosphorylation levels of JAK2 and STAT3 in the 22RV1 cell line. The selection of the 22RV1 cell line was driven by existing literature that reports the presence of PRLR, a factor deemed essential for delving into the molecular response to Osthole in prostate cancer[12]. Western blot analysis revealed that, in contrast to the untreated control group, the protein expression level of PRLR in 22RV1 cells was downregulated after a 48-hour exposure to Osthole. Simultaneously, there was a notable reduction in the phosphorylation of JAK2 and STAT3 (Figure 8). These observations align with the insights derived from network pharmacology, reinforcing the hypothesis that Osthole potentially combats prostate cancer by modulating PRLR, thereby inhibiting activation of the JAK2/STAT3 signaling cascade.

**Fig 8.**
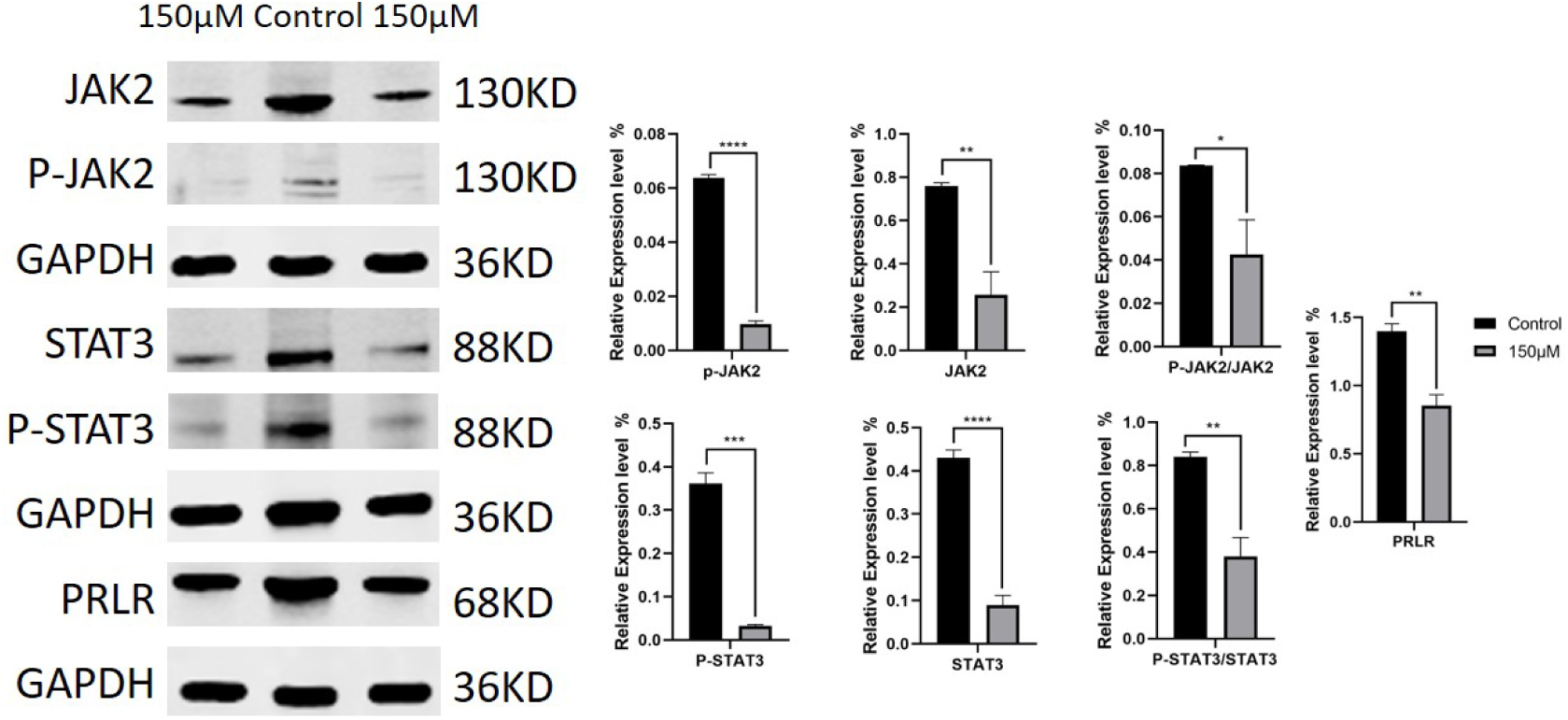
Western Blot Analysis. 150 μM Osthole significantly reduced the protein expression levels of PRLR, JAK2, and STAT3, as well as the phosphorylated forms of JAK2 and STAT3, and the ratio of phosphorylated to total JAK2 and STAT3 in 22RV1 cells.

## 4. Discussion

Our findings indicate that Osthole markedly suppresses the proliferation and metastasis of prostate cancer cells, triggers apoptosis, and attenuates tumor growth in in vivo models. Prostate cancer ranks among the leading malignant neoplasms in the male population worldwide, with an increasing trend in incidence and mortality observed in various geographic locations. Although current treatment strategies such as surgical intervention, radiation therapy, androgen deprivation therapy, and chemotherapy provide benefits for certain individuals, they are frequently associated with considerable side effects and show restricted effectiveness for late-stage or metastatic disease[13]. Therefore, the development of innovative pharmaceuticals is essential to bolster the therapeutic efficacy and elevate the living standards of affected individuals. Osthole, identified as a naturally occurring compound from a range of botanical sources, has recently demonstrated promising therapeutic potential in cancer research. It has been shown to significantly inhibit the growth of multiple cancer cell types, notably those associated with gastric and bladder cancer. Owing to its diverse biological functions, including anti-inflammatory, antioxidant, and anti-neoplastic capabilities, Osthole is a promising candidate for the development of new prostate cancer treatments[14].

This article initially explores the potential of Osthole to inhibit prostate cancer through network pharmacology analysis, utilizing multiple public databases to investigate possible targets and signaling pathways.By mapping the interactive network of the compound with biological entities, network pharmacology offers an encompassing view of Osthole’s molecular targets, its associated signaling cascades, and their intricate relationships within the biological context. This comprehensive method of research not only improves the accuracy and efficiency of drug development but also provides scientific evidence for the clinical application of the drug. We have identified targets such as AKT1, TNF, IL6, STAT3, and CTNNB1 in Osthole’s inhibition of prostate cancer, along with pathways including the TNF signaling pathway, the Interleukin-6-17 (IL-6-17) signaling pathway, and the prolactin signaling pathway. Studies have indicated that Osthole can inhibit prostate cancer by suppressing the TGF-β/Akt/MAPK pathway[8], which is consistent with our network pharmacology findings. However, our research also revealed that Osthole can exert anti-prostate cancer effects by binding to PRLR, thereby inhibiting the JAK2/STAT3 pathway. Prolactin (PRL), a polypeptide hormone secreted by the anterior pituitary gland in mammals, is a member of the growth hormone family. Its primary physiological role is to stimulate the growth and development of mammary glands and regulate milk production during late pregnancy and lactation[15]. In addition, PRL is involved in the regulation of the reproductive system, immune system, and various other physiological processes in the body[16]. Studies have indicated that PRL is a major contributor to the development of breast cancer and is upregulated in numerous hormone-dependent cancers, including prostate, ovarian, and endometrial cancers[17–19].

Based on the network pharmacology forecast, we verified the effects of Osthole on prostate cancer cell proliferation through cellular experimentation. By leveraging CCK8 and Transwell invasion assays for cell viability assessment, we established the growth-inhibiting role of Osthole in bladder cancer cells. While earlier research employed DU145 and PC3 human bladder cancer cell lines as models to study the suppressive effects of Osthole on prostate cancer[8], our study examined a more extensive range of prostate cancer cell lines. The experimental outcomes with the DU145 and PC3 cell lines were consistent with the findings reported in the aforementioned literature. However, the effects of Osthole on the 22RV1 cell line have not been documented in the existing literature to date. Therefore, our experiments have broadened the horizons of research in this area, and we observed that cells treated with Osthole exhibited nuclei that were smaller in size, deeply stained, and disorganized in arrangement, suggesting their potential pro-apoptotic capabilities.

Molecular docking studies were conducted to elucidate the molecular mechanisms underlying the action of Osthole. The results demonstrated robust binding interactions between Osthole and both the PRLR and STAT3 proteins, suggesting a theoretical framework whereby Osthole could potentially suppress the JAK2/STAT3 signaling pathway through its interaction with PRLR. This suggests that Osthole possesses the ability to inhibit both the proliferation and metastatic capabilities of breast cancer cells, primarily through the suppression of the JAK2/STAT3 signaling cascade[20]. However, the analogous potential of Osthole to exert inhibitory effects in prostate cancer, along with the intricate mechanisms driving this process, remains to be more rigorously explored and empirically confirmed.

In this study, we analyzed RNA sequencing data of prostate cancer patients from the TCGA database and identified that high PRLR expression was significantly associated with poor overall survival, suggesting its potential as a prognostic biomarker for prostate cancer. Previous studies have indicated that the association between PRLR and inflammatory response may be a key mechanism through which PRLR contributes to tumor progression. Chronic inflammation, as a critical component of the tumor microenvironment, can promote tumor progression by enhancing angiogenesis, immune evasion, and cell proliferation [21]. Therefore, PRLR may influence patient prognosis by activating inflammation-related pathways that reshape the tumor microenvironment.

Although JAK2 and STAT3 expression levels were not significantly correlated with prognosis in this study, it is noteworthy that these two genes have been implicated in tumor microenvironment regulation in other cancer types. For instance, JAK2 has been associated with tumor mutational burden in prostate cancer [22], and STAT3 is recognized as a pivotal regulator in inflammatory and tumor signaling networks [23]. Thus, JAK2 and STAT3 may not directly contribute to prostate cancer progression but could modulate the immune tumor microenvironment, thereby promoting disease development. Further analysis revealed significant enrichment of PRLR, JAK2, and STAT3 in the inflammatory response pathway, indicating that inflammation may serve as a central pathway mediating the combined effects of these genes. The inflammatory response pathway not only directly impacts cancer cell proliferation and survival but also promotes epithelial-mesenchymal transition (EMT) and immune suppression, driving tumor progression [24]. From a clinical perspective, the prognostic potential of PRLR provides a promising tool for risk stratification and personalized treatment of prostate cancer patients. Furthermore, the shared enrichment of PRLR, JAK2, and STAT3 in the inflammatory response pathway suggests that targeting this pathway could be a viable therapeutic strategy to inhibit their combined effects. For example, anti-inflammatory therapies could simultaneously mitigate PRLR’s role in inflammatory responses and suppress the deterioration of the tumor microenvironment.

To substantiate our hypothesis empirically, we performed western blot analysis, which demonstrated a marked reduction in the expression levels of PRLR in the 22RV1 prostate cancer cell line following treatment with Osthole. This significant downregulation of the PRLR protein indicates that Osthole may modulate the expression of key receptors involved in prostate cancer progression. Subsequent treatment with Osthole resulted in a notable reduction in the phosphorylation status of both JAK2 and STAT3 proteins, implying an inhibitory effect on the JAK2/STAT3 signaling pathway.Studies have shown that the expression of PRLR is not ubiquitous among prostate cancer cell lines[25], implying that the anticancer action of Osthole could be mediated through alternative pathways in addition to PRLR.The JAK2/STAT3 pathway is a prevalent signaling cascade that facilitates the transduction of signals from the extracellular milieu to the nucleus upon the binding of cytokines and growth factors to specific cell surface receptors[26]. Under normal conditions, the pathway is activated when prolactin, growth factors such as EGF, and cytokines, predominantly of the IL-6 family, bind to the extracellular domains of their respective receptor tyrosine kinases (RTKs)[27]. This signaling pathway is tightly regulated and serves a multitude of biological functions, including cell proliferation, differentiation, and apoptosis[28]. However, aberrant activation of the JAK2/STAT3 pathway is frequently detected in various tumors and has been implicated in the genesis, angiogenesis, and metastasis of many cancers[29]. Additionally, the JAK2/STAT3 pathway can interact with other pathways, such as the NF-κB and FOXO pathways, which are closely associated with cancer development [30–31].

In our detailed study of the effects of Osthole on prostate cancer, we found that it could suppress tumor growth and metastasis by targeting the JAK2/STAT3 pathway. This supports the accuracy of network pharmacology predictions and suggests the potential of Osthole as a treatment for prostate cancer. Further research could lead to more effective and safer therapies for such patients. Future research should further investigate the molecular mechanisms of PRLR within the inflammatory response pathway, with a particular focus on its potential synergistic interactions with JAK2 and STAT3. Although JAK2 and STAT3 did not show significant prognostic associations in the current analysis, functional experiments are warranted to validate their roles in inflammation-driven tumor progression. These findings could provide valuable insights into the development of anti-inflammatory strategies and targeted therapies, offering new directions for therapeutic intervention in prostate cancer.

In conclusion, this study validated the pharmacological mechanism of Osthole in prostate cancer using network pharmacology predictions and in vivo and in vitro studies. This suggests that Osthole inhibits the progression of prostate cancer by targeting the JAK2/STAT3 pathway. Therefore, Osthole is a potential drug for the treatment of prostate cancer.

## Author contributions

Linjun Yan: Writing – original draft, Resources, Investigation. Yuanqiao H: Conception and revision of the manuscript Qi Cui: Resource Investigation. Xiaohong Wang: Formal analysis and methodology. Feng Lv: Writing, review, and editing. Keyue Cao: Writing, review, and editing. Yuanjian Shao: Methodology, Visualization. Yuanjian Shao: Funding acquisition, Supervision, Project administration. All the authors contributed to the writing and review of the manuscript and approved the final version for submission.

## Ethical approval

Animal ethics: The experimental protocol for the mice used in the PDX models was approved by the IACUC of Royo Biotech Co., Ltd. (RYE2022080301, RYE2023112501 Nanchang, China). Animal transport, husbandry, and experimental procedures were performed in accordance with national legislation (SN/T 3986-2014) on the protection of animals used for scientific or educational purposes. After the surgical operations and experiments, the animals were euthanized by CO_2_ asphyxiation. All methods were performed in accordance with the ARRIVE guidelines for reporting animal experiments.

## Data Availability

Data is provided within the manuscript or supplementary information files.Allimages in this work are reproducible and may becopied for educational or research purposes, without requiring further permission from the authors or publishers.

## Consent for publication

All authors read and approved publication.

